# Comparative Analysis of Acoustic Propagation Parameters of Natural Sounds of *Anopheles gambiae s.s* and *Odorrana tormota* Significant in Mosquito Startle

**DOI:** 10.1101/725960

**Authors:** P. A. Mang’are, F. G. Ndiritu, S. K. Rotich, J. K. Makatiani, B. W. Rapando

**Affiliations:** Physics Department, Egerton University. P. O Box 536, Egerton, Njoro. Kenya; Physics Department, Masinde Muliro University of Science and Technology. P. O Box 190-50100, Kakamega. Kenya; Department of Physics and Mathematics., Moi University, P. O. Box 3900-30100 Eldoret. Kenya; Department of Biological Sciences., Moi University, P. O. Box 3900-30100 Eldoret. Kenya

**Keywords:** Vector, Pathogens, Acoustic Energy, Fundamental frequencies, Harmonics

## Abstract

Acoustics of varied frequency ranges generated naturally by animals or artificially by electronic devices have shown startle effect to insects. It has been shown that mosquitoes use the reactive near-field in antennae communication with negative phonotaxis in male *Aedes diantaeus* evoked by low frequency acoustic signals of a carrier frequency 140–200 Hz. Also, studies with the 35-60 kHz *Odorrana tormota* sound recorded a 46 % repellence in female *Anopheles gambiae*, the malaria vectors. Declining malaria morbidity and mortality is attributed to current vector and pathogen interventions. However, the rate of decline in malaria morbidity and mortality is impeded by buildup of resistance in pathogens and vectors to chemicals. This study therefore characterised animal sounds essential for further investigation in the control of malaria through mosquito startle. The research determined, analysed and compared the acoustic propagation parameters of the recorded natural sounds of the male *Anopheles gambiae*, female *Anopheles gambiae* and *Odorrana tormota* using Avisoft SASLAB Pro and Raven Pro 1.5. All sounds were observed to have frequency modulation with harmonics stretching to ultrasonic levels. Uniquesly, the sound of *O. tormota* showed constant frequency modulation. The pupae of *A. gambiae* were reared in vials quarter filled with water and covered with a net at 60-80 % humidity, 25±2 °C temperature and equal light-darkness hour cycle at Kenya Medical Research Institute (KEMRI) entomology laboratories. The parameters showed a significant deference in fundamental frequency (maximum entire), Peak amplitude (maximum), peak amplitude (mean), Peak amplitude (mean entire) and peak amplitude (maximum entire) of the sound of male *A. gambiae* and *O. tormota* (p < 0.05). The maximum frequency (minimum entire) of both sexes of *A. gambiae* was equal (1.90 kHz) with variability being observed in maximum frequency (end), maximum frequency (maximum), maximum frequency (mean), maximum frequency (maximum entire) and maximum frequency (mean entire). Frequency (maximum). A paired samples t-test comparison of the maximum frequency (mean), maximum frequency (maximum), maximum frequency (end), maximum frequency (maximum entire) and maximum frequency (mean entire) of the sound of the female *A. gambiae* and male *A. gambiae* indicated no significant difference between the sounds (p > 0.05). The maximum frequency (mean) of the sounds of both sexes of *A. gambiae* correlated highly negative (r = −0.658). The bandwidth (end), bandwidth (maximum), bandwidth (maximum entire), peak amplitude (mean) and bandwidth (mean entire) of the sound of the male compared with female *A. gambiae* differed significantly. The signal power for the non-pulsate sounds of the male *A. gambiae* remained almost constant at 80 dB from 10 kHz to 65 kHz beyond which the acoustic energy declining to 45 dB. Also, the sounds of the female *A. gambiae* did not exhibit any spikes in power but remained steady at 85 dB from 10 kHz up to 60 kHz beyond which the acoustic energy declined to 50 dB. The signal power of the pulsate sound of *O. tormota* was 89 dB. The propagation parameters of the male mosquito and O. tormota compared favourably indicating its potential in the startle of the female mosquito.

**The author summary:** Philip Amuyunzu Mang’are is a PhD. Physics student in Egerton University. He has authored many papers and books. He is currently a Lecturer of Physics (Electronics), Masinde Muliro University of Science and Technology. He is a member of the Biophysical Society and the current President of Biophysical society (Kenya). Prof. Ndiritu F. Gichuki, is a Professor of Physics Egerton University. Currently he is the Registrar Academic Affairs in Chuka University. His vast experience has seen him supervise many postgraduate students who have taken key positions in the society. Prof. Samwel Rotich is a Profesor of Physics in Moi University specialising in Electronics. He has a wide experience in Physics and Biophysics. He is a registered member of the Biophysical Society and the Patron of Biophysical Society Kenya Chapter. He has published many papers and supervised many postgraduate students. Dr. Makatiani Kubochi is a Lecturer in Moi University with vast experience in entomology. She has published many papers and supervised many postgraduate students. Dr. Rapando Bernard Wakhu is a renown theoretical Physicist with experience in acoustics and Fourier analysis based in Masinde Muliro University of Science and Technology. He has supervised many postgraduate students and published many papers.

## 1. Introduction

The *Anopheles gambiae* and the *Odorrana tormota* generate sounds naturally whose frequencies stretch to ultrasonic levels. The focus of this study was to comparatively analyse the acoustic propagation parameters of natural sounds of *A. gambiae* and *O. tormota.* Optimal parameters determined from this study were essential in the investigation of the startle responsiveness of the African female *A. gambiae* to ultrasound and the design of a mosquito repellent.

### 1.1 The biology of the *Anopheles gambiae* and *Odorrana tormota*

Mosquitoes have four distinct stages in their life cycle which are recognized by their unique appearance; which include the egg, larva, pupa, and adult (CDC, 2010). The egg, larva and pupa are aquatic stages in the lifecycle of the *A. gambiae* and last 5-14 days, depending on the species and the ambient temperature (CDC, 2010). The non-drought resistant eggs having floats are between 0.47mm and 0.48 mm long, convex below and concave above, with the surface covered with a polygonal pattern (Gillies and de Meillon 1968; Foster and Walker 2009; CDC 2010). The organic matter and algae feeding larvae ranges between 5mm to 6 mm in size and lie parallel to the water surface in order to breathe due to lack of respiratory siphons (Gillies and de Meillon, 1968; Garros *et al*., 2008; Foster and Walker, 2009). The non-feeding pupae are comma-shaped when viewed from the side and are very mobile (Foster and Walker 2009; CDC 2010). Adult female *Anopheles* have palps which are as long as their proboscis, an average wing length varying from 2.8 to 4.4 mm and rest with their abdomen raised into the air (Gillies and de Meillon, 1968; Foster and Walker, 2009). The adult mosquitoes have slender bodies consisting of the head, thorax and abdomen; the head specialized for acquiring sensory information and for feeding. The mosquito antennae detects host and breeding sites odors (CDC, 2010). The head also has an elongated, forward-projecting proboscis used for feeding, and two sensory palps. The adult stages of many mosquito species are feeders of blood, which has given some disease causing organisms a reliable mode of transmission to animal hosts. It is during the adult stage that the female *Anopheles* mosquito acts as malaria vector. The adult females can live up to a month or more in captivity but they don’t live more than 1-2 weeks in nature (Antonelli *et al.*, 2007). Both male and female adult mosquitoes feed on plant nectar, but the female feed on vertebrates’ blood, for nutrients required for egg maturation (Foster and Walker, 2009). Female Anopheles mosquitoes lay eggs on the surface of the water at night and under favorable conditions, hatching occurs within one or two days and develop within the aquatic habitat. The *A. gambiae* larvae develop in permanent man-made structures and natural pools (Kweka *et al*., 2012). It is important to understand the lifecycle of the mosquito so as to effectively control them. Anthropophilic biting female mosquitoes not only irritate people and animals, but also transmit malaria. The body parts of the adult stage of the mosquitoes, mainly the antennae serve an important role in communication (Mohankumar, 2010).

The *Odorrana tormota* belongs to kingdom *Animalia*, Phylum *Chordata*, Class *Amphibia*, Order *Anura* and of the family *Ranidae*. The synonyms for *Odorrana tormota* are *Amolops tormotus*, *Odorrana tormotus* and *Rana tormotus* (Wu, 1977; Huiqing and Ermi, 2004; Frost; 2013). The *Odorrana tormota* shelters in moist rock crevices during the day, and is found in thick brush alongside streams at night (Fei, 1999). The reproductive season begins in June and they lay eggs which are milky yellow in color, and measure about 2.0 mm in diameter (Fei *et al*., 1999). The existence of tadpoles has not been recorded for this species (Huiqing and Ermi, 2004). The *Odorrana tormota* is a frog restricted to Huangshan in Anhui Province, and Jiande and Anji counties in Zhejiang Province, China generating ultrasounds through vocal apparati (Arch *et al*., 2008; Shen *et al*., 2011). The frog uses the frequency range of up to 128 kHz for communication (Arch *et al*., 2008). During the reproductive season, males emit a variety of high-pitched calls at night with energy spectrums extending into the ultrasonic range (Feng *et al*., 2006; Feng *et al*., 2009). Recent research with the *O. tormota* calls showed some degree of downward frequency modulation with a subset of calls having a carrier of constant frequency (Feng *et al*., 2002; Arch *et al*., 2008). As observed in recent studies, ultrasound from *O. tormota* can play a critical role in malaria vector control by evoking startle responses in malaria vectors since the call frequencies stretch beyond the startle range of 38 – 44 kHz in mosquitoes (Mohankumar, 2010; Mang’are *et al*., 2012). The sound of *O. tormota* having exhibited the greatest startle responses in mosquitoes needs to be further investigated besides the sounds of the male *Anopheles gambiae* (Mang’are *et al*., 2012).

### 1.2 Mosquito audition, communication and antennae theory

The mosquito has a pair of large, wraparound eyes, and a pair of long, hairy antennae; its ears projecting from the front of its face (Hoy, 2006). The antenna detects the particle velocity component of a sound field, which is restricted to the immediate vicinity of the sound source in acoustic near field. Ultrasound generated artificially or naturally is detected by mosquitoes evoking evasive response (Mohankumar, 2010). The mosquito flight tone is an unusual communication signal in that its production is directly linked to locomotion, only varying the carrier frequency (Arthur *et al*., 2014). The flight tone is characteristic of a species and can sometimes be used to identify species or count individuals (Raman *et al*., 2007). Mosquitoes use flight tones as communication signal with the male mosquitoes being attracted to real and artificial female flight tones (Wishart and Riordan, 1959).

The mosquito antennae consists of an elongated flagellum and a chordotonal organ housed within the pedicel at the base of each antenna called the Johnston’s organ (Gibson *et al*., 2010). The flagellum vibrates when stimulated by periodic air displacements (Göpfert and Robert, 2000). At its base, the flagellum tapers into a structure of 60–80 prongs which resemble the spokes of an upturned umbrella (Gibson *et al*., 2010). The prongs are attached to many thousands of mechanosensory sensillae known as the scolopidia of the Johnston’s organ.

Insect hearing organs are divided tympanal ears, which are sensitive to the pressure component of sound, and movement receivers, which respond to oscillations of air particles in the sound field (Hoy and Robert, 1996). Movement receivers are usually light structures such as sensory hairs or antennae that are deflected by the sound-induced oscillations of air particles. The antennae are sexually dimorphic, with males having a greater number of sensillae in the Johnston’s organ (∼14,000 in males, ∼7,000 in females) and the greater number of fibrilla on the flagellum led to a conclusion that mating behavior involved audition (Gibson and Russell, 2006). The greater surface area of the male flagellum increases their responsiveness to particle displacement (Göpfert *et al*., 1999).

Mosquitoes are affected by ultrasounds in the range 38 kHz-44 kHz (Mohankumar, 2010). Ultrasound creates stress on the nervous system of mosquito jamming its own ultrasound frequency immobilizing it. Ultrasound also initiates avoidance behavioural response from the source (Mohankumar, 2010). The male’s auditory system is selectively tuned to female flight frequencies of approximately 300Hz – 400Hz with maximum intensity at 380 Hz for *A. gambiae* (Gibson and Russell, 2006). The sound is transmitted in air as frequency modulated wave (FM) and activates the antennae (Maweu *et al.*, 2009). The female mosquitoes which detect the presence of male mosquitoes by sensing the 38 kHz ultrasound require blood meal for egg maturity after breeding (Baldini *et al.*, 2013; Imam *et al.*, 2014). The bred Female *Anopheles gambiae* become refractory to the males to avoid further breeding. The antenna, with its fine senses movements of air particles impinging acoustic waves that excite the sensory receptors contained within the JOs to generate action potentials that flood into the insect’s brain (Göpfert and Robert, 2000; Hoy, 2006). The sensillae mechanoelectrically transduce and amplify the nanometer sound-induced vibrations into electrical signals (Göpfert and Robert, 2000).

Experiments with recorded flight tones of mosquitoes showed that the individual male and female mosquitoes flew at mean fundamental wing-beat frequencies with males flying at significantly higher frequencies (∼700 Hz) than their conspecific females (∼460 Hz) (Gibson *et al*., 2010). The auditory range was noted to be ∼2 cm (Pennetier *et al*., 2010). The less sensitive female’s Johnston’s organ, compared to males, responds to antennal deflections of ± 0.00058 induced by ± 11nm air particle displacements in the sound field which surpasses other insects’ sensitivity (Göpfert and Robert, 2000). Ultrasound causes neural stress to mosquitoes evoking a startle response. Mosquitoes like many other insects avoid bat ultrasonic sound that the electronic mosquito repellent devices imitate (Mohankumar, 2010).

The focus of this study is to analyse the acoustic propagation parameter of the sounds of the *A. gambiae* and *O. tormota* essential for further study on use the responsiveness of the African female *A. gambiae* to the sound.

Electromagnetic communication between insects has also been observed with the antennae playing the role of the receiver or transmitter (Abdolali *et al*., 2013). Many insects use antennal or cercal structures to detect air particle vibrations from near-field sound sources (Sane and McHenry, 2009). The antenna theory enhances the understanding of communication in mosquitoes (Maweu *et al*., 2009). The hairs on the mosquito antennae serve as the dipole lengths and are equally spaced, with the dipole lengths range from 0.05 to 0.14 cm for the *A. gambiae, Culex pipiens* and *Aedes aegypti*. It was established that there were 20 dipoles stemming from the 0.26 cm long transmission line (Maweu *et al*., 2009). Basically, mosquitoes use the reactive near-field in antennae communication.

### 1.3 Mosquito mating behaviour

Male mosquitoes require about 24 hours before their *terminalia* get rotated and their fibrillae mature enough to become erect and detect females whereas the female mosquitoes need 48-72 hours before they become receptive to males prior to blood feeding in the wild (Clements, 1992). *Anopheles* males can mate several times, but females become refractory to re-insemination and re-mating is rare (Mohankumar, 2010). Mating in *anopheline* mosquitoes occur during the early evening, primarily in swarms (Diabaté *et al.*, 2011). The swarming males use their erect antennal fibrillae to detect a nearby female mosquito’s wing beat frequencies (Clements, 1992). The audible frequencies received by the male antennae ranges from 150 Hz to 500 Hz and evoking sexual behaviour that involves shifting pitches of their buzzes until they synchronize. Same sex mosquitoes’ sounds cannot converge (Nijhout and Craig, 1971). The case study on *Toxorhynchites* showed that the males harmonized their wing beat with females as they neared, for species recognition, before mating commenced (Gibson and Russell, 2006). Flying mated female mosquitoes produce familiar whining sound when searching for proteins (Maweu *et al*., 2009). This sound is very critical in this study as an audio switching system.

### 1.4 Electronic mosquito repellents (EMRs)

Electronic mosquito repellents (EMRs) are designed to repel female mosquitoes by emitting high-pitched sounds (Enayati *et al*., 2010; Ashim *et al.*, 2017). There exist many electronic mosquito repellents on the market which manufacturers argue that they are effective in repelling or attracting mosquitoes, though not common in Africa. The Anti-Pic®, Mosquito Repeller® DX-600 and Bye-Bye Mosquito® electronic mosquito repellents have been studied in order to establish their effectiveness in mosquito repellence but none has supported the claims of their effectiveness in mosquito repellence (Andrade and Bueno, 2001). The electronic mosquito repellents that generates ultrasound at 38 kHz imitating the male mosquitoes have been designed. It was also noted that in 12 of the 15 experiments, the landing rates of mosquitoes on the human participants in the groups with functioning EMR was actually higher than in the control groups (Enayati *et al*, 2010).

Ultrasound causes nervous stress to the mosquito and at the same time evoke fear due to predation or/and further mating depending on the source of the natural ultrasound (Mohankumar, 2010). Recent research findings with recorded natural ultrasound showed evidence of startle responses in mosquitoes (Enayati *et al*, 2010; Mohankumar, 2010; Mang’are *et al*., 2012). The studies about the repellence of the female *A. gambiae* using the hearing mechanism is still a viable venture based on the confirmation that in both males and females, the antennae are resonantly tuned mechanical systems that move as simple forced damped harmonic oscillators when acoustically stimulated, despite the differences in the hearing ability (Göpfert *et al*., 1999). Recent experiments demonstrated that tonal acoustic signals with a carrier frequency of 140–200 Hz yielded avoidance responses (negative phonotaxis) of male *Aedes diantaeus* swarming mosquitoes (Culicidae) in the range of 140– 200 Hz (Lapshin and Vorontsov, 2018).

### 1.5 Statement of the Problem

High mortality and morbidity in Africa and the world at large is caused by malaria whose vectors are the mated female *A. gambiae*. Malaria has also caused a large economic burden to both individuals and countries in an effort to control the vectors or treat malaria. There is a slowed achievement of satisfactory reduction levels in mortality and morbidity due to buildup of resistance in chemicals used to treat malaria or control the vectors. Complimentary approaches that target the vectors have been recommended. Animal sound mimicking electronic mosquito repellent devices (EMR) have been used though yielding paltry 20 % repellence. Recorded 35 – 60 kHz ultrasound from *O. tormota* showed improvement (46 % repellence) in evasive response by the mated female *A. gambiae*. It is on this premise that an investigation into ultrasound from the male *A. gambiae* was investigated. Also the sound of the female *A. gambiae* was investigated in order to obtain the characteristic frequencies. Recent researches have shown refractory behaviour of the mated female *A. gambiae* to the male *A. gambiae* through detection of their ultrasound using the antennae. This research focused on the comparative analysis of the acoustic propagation parameters of the natural sounds of male and female *A. gambiae* and *Odorrana tormota.* The research determined the acoustic propagation parameters of the sounds of the male and female *A. gambiae*, analysed and compared the acoustic propagation parameters of the sounds of the male *A. gambiae* and *O. tormota.* The parameters of the sound of male and female *A. gambiae* were compared with those of the sounds of *O. tormota* in order to determine superior parameters essential for the design of a mosquito repellent device, an additional tool in mosquito repellence.

### 1.6. Objectives

#### 1.6.1. General Objective

Analysis and comparison of the acoustic propagation parameters of the natural sounds of the male and female *Anopheles gambiae s.s* and *Odorrana tormota*

#### 1.6.2. Specific Objectives

i. To determine the acoustic propagation parameters of the sounds of the male and female *A. gambiae*
ii. To analyse and compare the acoustic propagation parameters of the sounds of the *A. gambiae s.s* and *O. tormota*

## 2. Materials and Methods

### 2.1. Sound of *A. gambiae s.s* and *O. tormota*

The pupae of *A. gambiae* were reared in vials quarter filled with water and covered with a net at 60-80 % humidity, 25±2 °C temperature and equal light-darkness hour cycle at KEMRI, Entomology laboratories. The brightness in the laboratory was improved by electric light from fluorescent tubes aerially located for uniform supply. The sounds of the *A. gambiae* and the Chinese frog, *O. tormota* were investigated. A computer mounted with a hardlock key to run of Avisoft-SAS LAB Pro version 5.2 programme and a recorder was used for recording the sounds of the male and female *A. gambiae*. The male *A. gambiae* were identified based on their many flagella (fine hairs) on the antennae, noticeably bushy to the eye. The size of the male *A. gambiae* is relatively smaller compared to the female *A. gambiae*. Practically the males were noted not to feed on blood meal. The female have sharp mouthparts, bigger in size, emitting the annoying buzzy sound and fed on blood meal. The recorder consisted of the Avisoft UltraSoundGate (model 112) connected to an omnidirectional microphone, set to default and connected to the computer through the universal serial bus (USB) port. The Avisoft-SAS LAB Pro, version 5.2 software was open and the microphone directed to the source of sound. With the gain on the Avisoft ultrasound Gate (model 112) adjusted to an appropriate level (50 %) to avoid over modulation and the recording level from the computer set to 20 dB, the recording button was pressed to record the sound. The sound was recorded from a swarm of 100 mated, 3 days old male and female *A. gambiae* at a sampling frequency of 500 kHz at 16 bit from a cylindrical glass cage covered at both ends with a netting as given in Figure 1. An aspirator was used to transfer mosquitoes from the resident cage to the cylindrical cage for sound recording.

**Figure 1:**
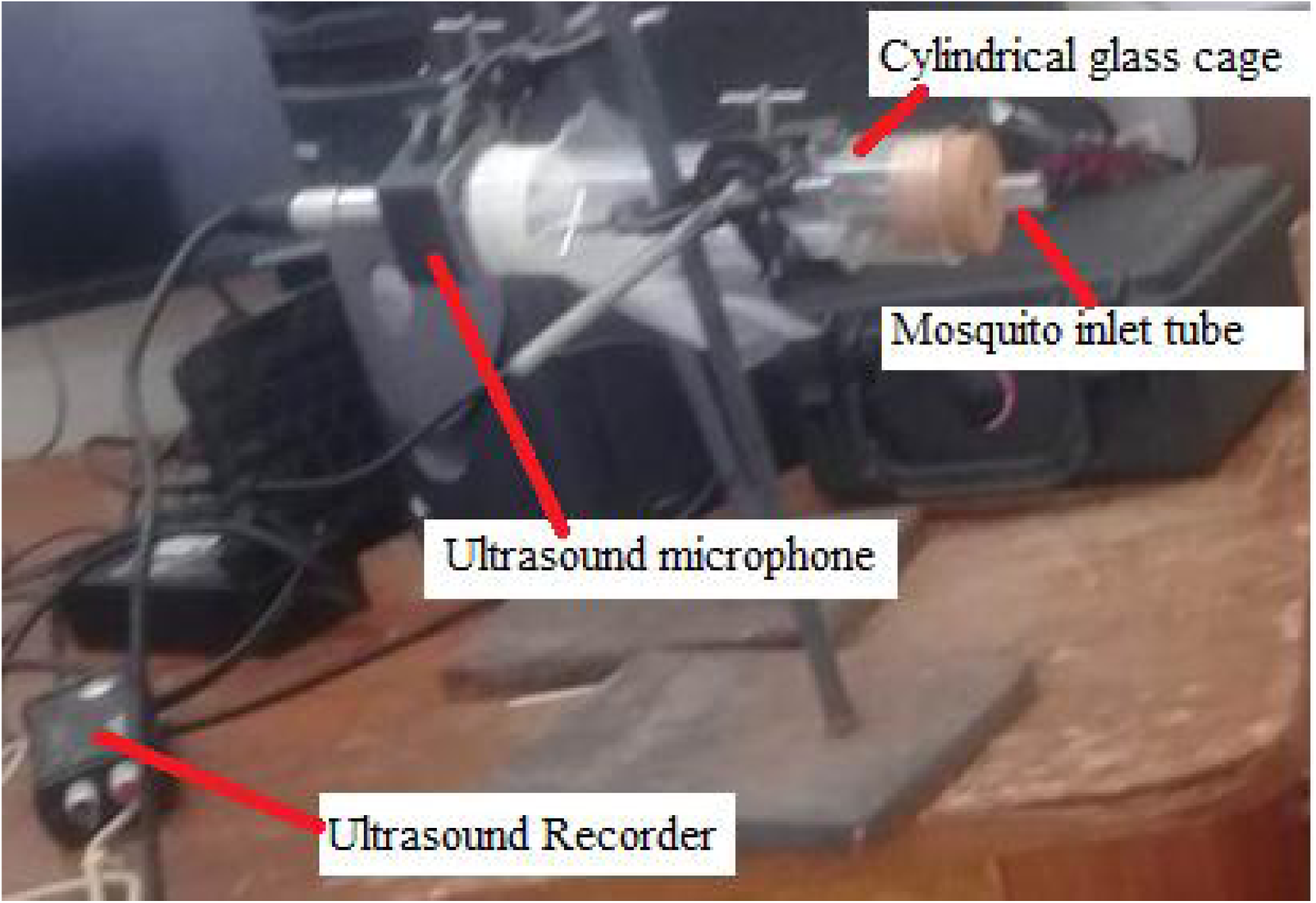
The Sound recording setup

The sound of *O. tormota* was recorded by 702 digital recorder from the Huangshan Hot Springs, Anhui Province in China at a sampling frequency of 192 kHz at 16 bit. The sounds were acquired from Prof. Feng formerly of Illinois University. The Avisoft-SAS LAB Pro version 5.2 and Raven Pro 1.5 software was used for conversion of the sampling frequency of the sound of *O. tormota* from 192 kHz to 500 kHz and for compatibility hence allowing for analysis.

### 2.2. The acoustic propagation parameters of sounds of male and female *Anopheles gambiae s.s*

The sound of both male and female *Anopheles gambiae*; and the sound of *O. tormota* were subjected to Fourier transforms and analysis for determination of acoustic propagation parameters. This was achieved through automatic parameter measurements. The following settings were made to the analysis softwares: from the tools option, the calibration was set to SPL with reference to sound and the SPL reference was 20µPa which is the threshold. For the parameter generation which include amplitude and energy, the calibration method was set to SPL with reference sound for Channel 1and at a /gain (dB) set to zero under the tools menu. The full-scale range was calibrated to 19.8936 Pa and 17.6345 Pa for the male and female *A. gambiae* respectively. The envelope was set to original waveform whereas the pulse detection was set to gate function. The Fast Fourier transform (FFT), an option under the spectrogram parameters was set to 512 and hamming window selected for the display. The temporal resolution overlap was set to 50% with the colour palette set to graypal. The frame size was set to 100% for real time spectrogram parameters and the black and white box (B/W) checked for display.

The parameters determined using Avisoft SASLab Pro version 5.2 and Raven Pro. 1.5 included call duration, acoustic energy, peak frequency (mean), peak amplitude (mean), minimum frequency, maximum frequency (mean), bandwidth (mean), peak frequency (minimum entire), minimum frequency (minimum entire), maximum frequency (minimum entire), bandwidth (minimum entire), peak frequency (maximum entire), peak amplitude (maximum entire), minimum frequency (maximum entire), maximum frequency (maximum entire), peak frequency (mean entire), peak amplitude (mean entire), minimum frequency (mean entire), and maximum frequency (mean entire). The acoustic energy whose SI unit is Pa^2^s is a product of the square of amplitudes and sample time. The energy produced is the sum of the squared amplitudes multiplied by time. Also, 1 Pascal pressure is equal to SPL of 94 dB. The data obtained was transferred into an excel worksheet for editing and further analysis through the direct data exchange (DDE)/ Logfile settings. The acoustic propagation parameters of the natural sounds were analysed statistically using the same softwares and SPSS.

## 3. Results and Discussion

### 3.1. Determination of Transmission Parameters of Sounds of *Anopheles gambiae*

#### 3.1.1. Generation, Modulation and Spectral features of the Sounds of *A. gambiae* and *O. tormota*

A total of 1,496 and 1,161 calls of the male and female *A. gambiae* respectively generated naturally through wing-beat were studied. The calls of the male *A. gambiae* were compared to the 684 calls of *O. tormota* naturally generated through vocal apparati (Arch *et al*., 2008). The calls studied lasted for a minimum duration of 62.13 s and a maximum duration of 150.21 s for the female *A. gambiae*. The calls for the female *A. gambiae* took an average duration of 118.82 s. The calls for the male *A. gambiae* took a minimum duration of 26.54 s and a maximum duration of 139.72 s with an average duration of 85.09 s. The female *A. gambiae* were strong enough at three days to sustain longer duration of wing-beat compared to the male mosquitoes of the same age. The spectral features in the combined spectrograms shown in Figure 2 reveal Frequency modulation (FM) in both the male and female *A. gambiae* essential in communication (Wishart and Riordan, 1959; Raman *et al*., 2007; Maweu *et al*, 2009, Arthur *et al*., 2014). Large modulations in males could indicate fitness in terms of the ability to fly and maneuver at suboptimal wingbeat frequencies (Arthur *et al*., 2014).

**Figure 2:**
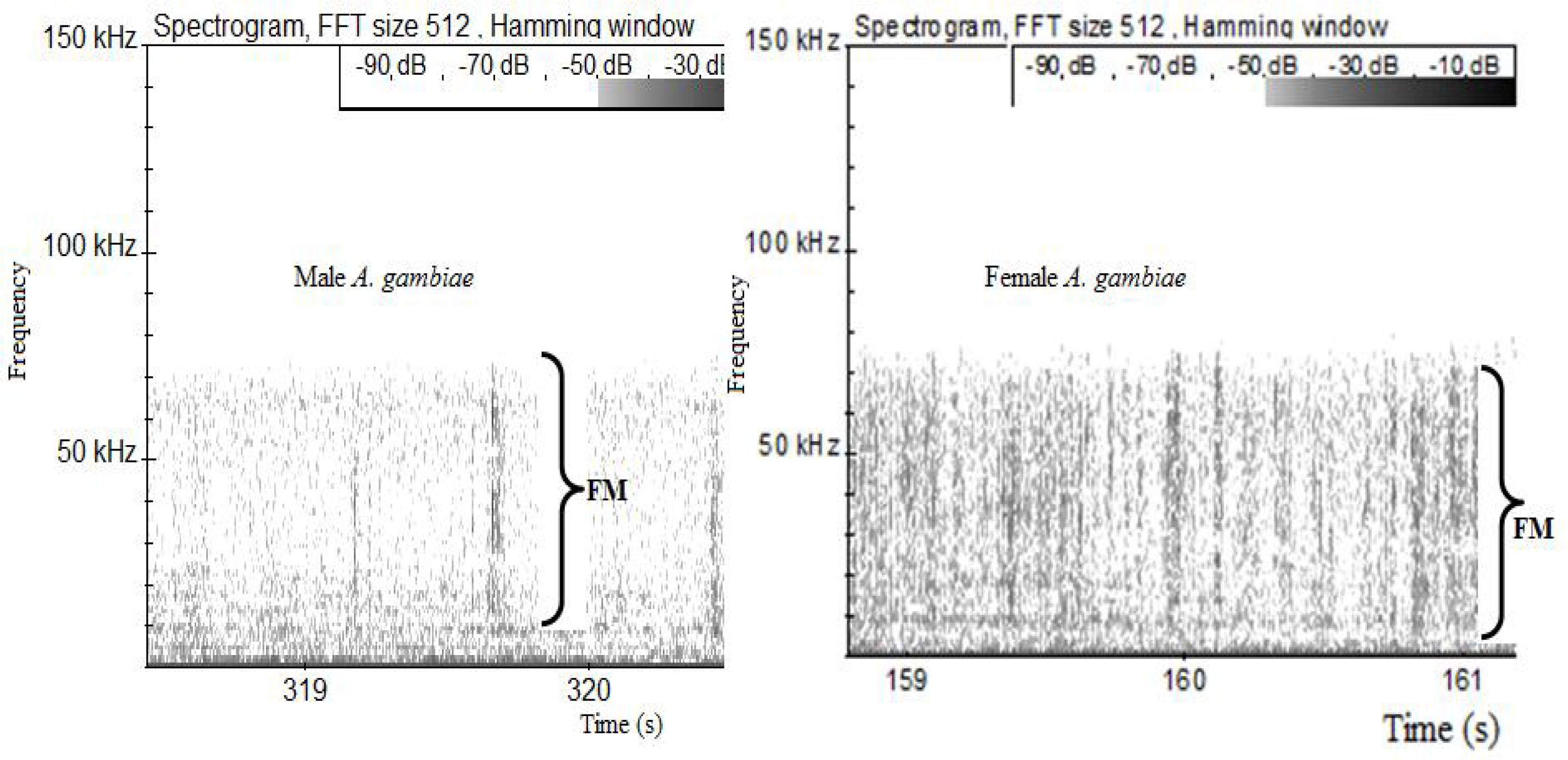
Modulation and spectral features in the sound of the *A. gambiae*

The calls from both male and female *A. gambiae* consists of several harmonic segments with energy extending into the ultrasonic range. The signals exhibit signal breaks and a chaotic nonlinear characteristics. The female *A. gambiae* was dominated by the chaotic nonlinear characteristic. The pulsate calls of the Chinese frog *O. tormota* had proven effective in the startle of the female *A. gambiae*, yielding 46 % repellence, evidence for the feasibility for using ultrasound in mosquito control (Mang’are *et al*., 2012). The sound of the male *A. gambiae* represented by the oscillogram in Figure 3a is minimally pulsate in nature though critical in investigation of mosquito startle.

**Figure 3a:**
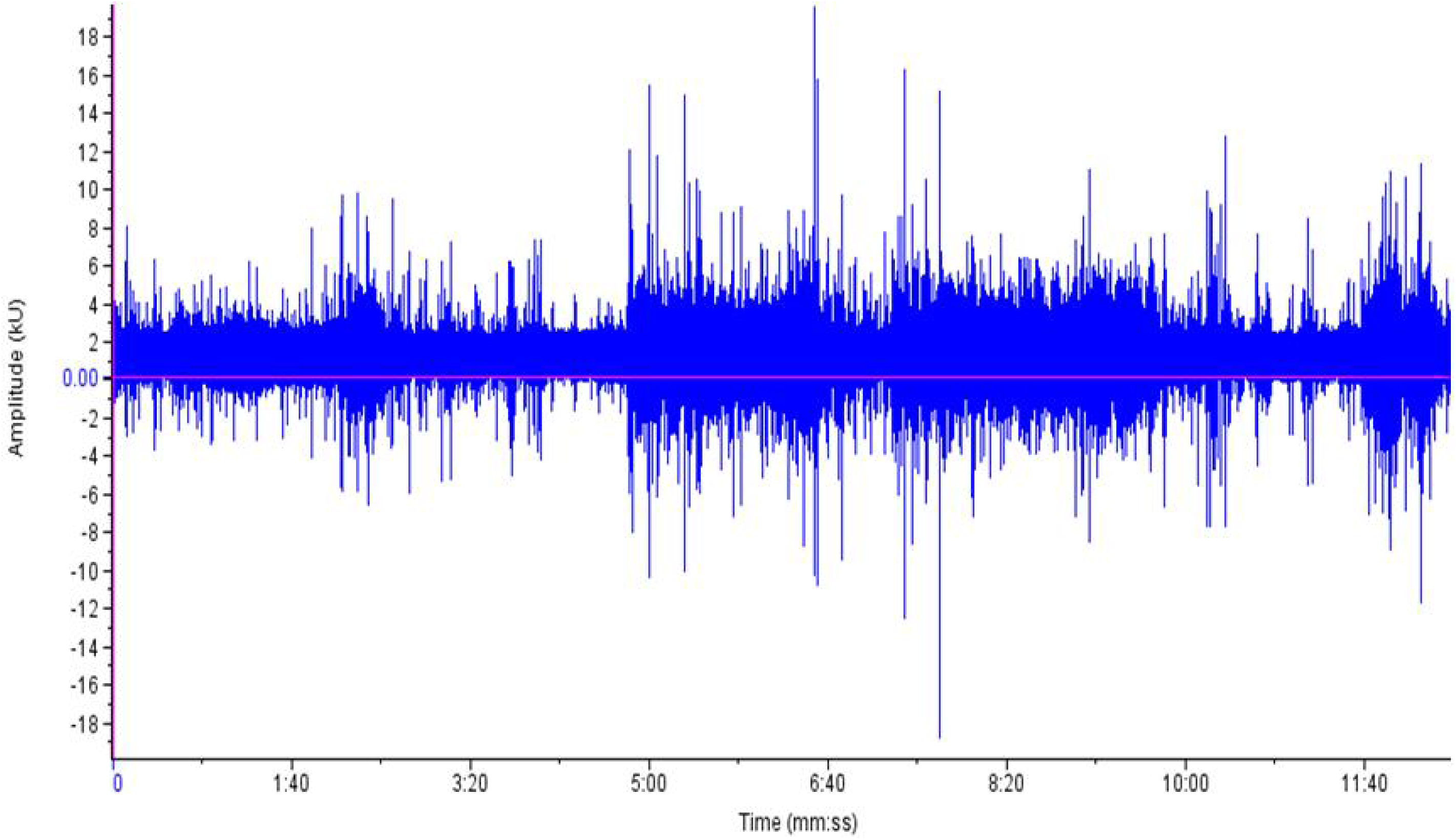
Oscillogram for the sound of male *A. gambiae*

**Figure 3b:**
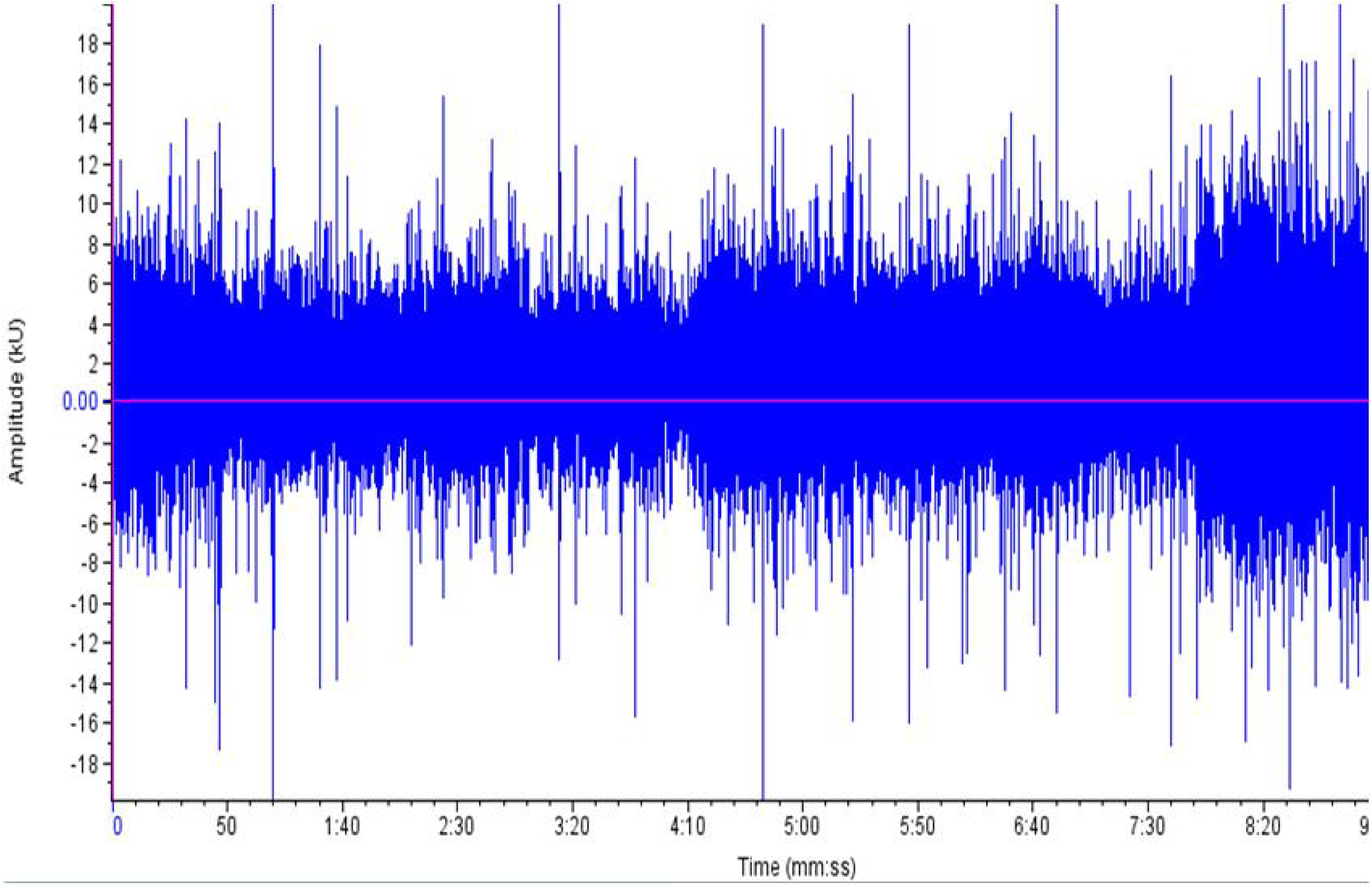
Oscillogram for the sound of female *A. gambiae*

Similarly, the sound of the female *A. gambiae* represented by the oscillogram in Figure 3b is minimally pulsate.

The sound of *O. tormota* is characterised by frequency modulation (FM) and constant frequency (CF) modulation as shown in Figure 4 as reported in recent findings (Feng *et al*., 2002; Arch *et al*., 2008). It is also characterised by harmonics and subharmonics nonlinear spectral features (Feng *et al.*, 2009). There also exists signal breaks in the sounds of male *A. gambiae*, female *A. gambiae* and the *O. tormota* as evidenced in the spectrogram in Figure 2 and 4 associated with change in locomotion.

**Figure 4:**
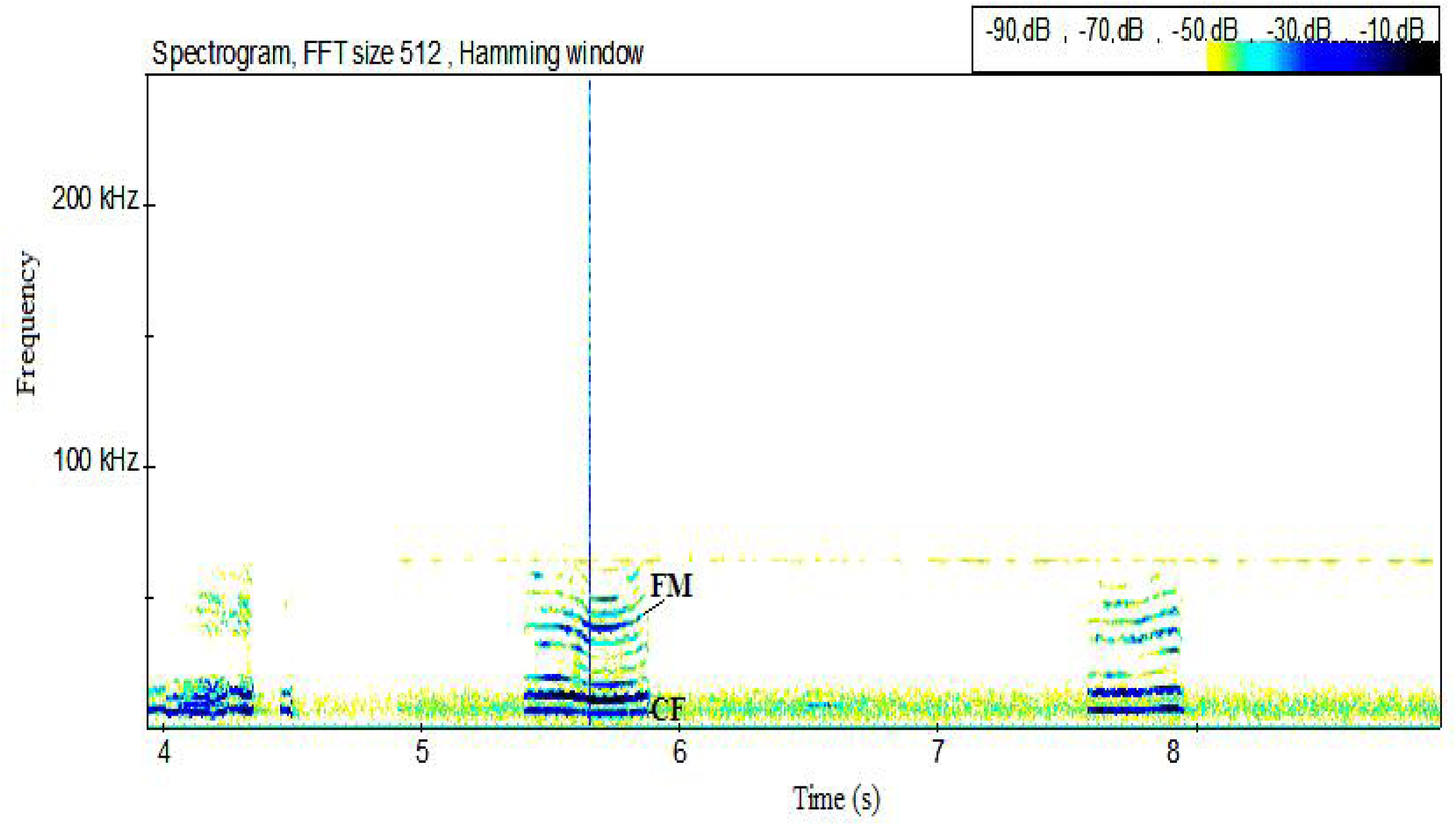
Spectral features in the sound of *O. tormota*

#### 3.1.2. Acoustic Transmission Parameters of the sounds of *A. gambiae*

The acoustic parameters were determined and analysed using the Avisoft SasLab Pro 5.2 and Raven Pro 1.5 softwares. The analysed data for the minimum value, maximum value and mean is given in Table 1, 4 and 5.

**Table 1:** Fundamental Frequencies and Harmonics

| Parameter (in Hz) | Female *A. gambiae* |  |  | Male *A. gambiae* |  |  |
| --- | --- | --- | --- | --- | --- | --- |
|  | Minimum | Maximum | Mean | Minimum | Maximum | Mean |
| Fundamental (end) | 0 | 900 | 180 | 0 | 1000 | 411.11 |
| Fundamental (maximum entire) | 2100 | 9100 | 3660 | 1000 | 2100 | 1466.66 |
| Fundamental (maximum) | 0 | 900 | 360 | 0 | 1000 | 411.11 |
| Fundamental (mean entire) | 1000 | 1000 | 1000 | 900 | 1000 | 955.56 |
| Fundamental (mean) | 900 | 900 | 900 | 900 | 900 | 900 |
| Fundamental (Minimum entire) | 900 | 900 | 900 | 900 | 900 | 900 |
| Fundamental (start) | 900 | 900 | 900 | 900 | 900 | 900 |
| Maximum Frequency (end) | 15600 | 77100 | 58760 | 9700 | 68300 | 40000 |
| Maximum Frequency (maximum) | 4800 | 72200 | 43500 | 4800 | 76100 | 48433.3 |
| Maximum Frequency (mean) | 20500 | 68300 | 45260 | 2900 | 25300 | 12322.2 |
| Maximum Frequency (maximum entire) | 77100 | 82000 | 80240 | 71200 | 81000 | 78066.6 |
| Maximum Frequency(mean entire) | 14900 | 58400 | 30240 | 2900 | 36300 | 21055.5 |
| Maximum Frequency(Minimum entire) | 1900 | 1900 | 1900 | 1900 | 1900 | 1900 |
| Minimum Frequency (end) | 900 | 900 | 900 | 900 | 900 | 900 |
| Minimum Frequency (maximum) | 900 | 900 | 900 | 900 | 900 | 900 |
| Minimum Frequency (mean) | 900 | 900 | 900 | 900 | 900 | 900 |
| Minimum Frequency (minimum entire) | 900 | 900 | 900 | 900 | 900 | 900 |
| Minimum Frequency(maximum entire) | 1900 | 1900 | 1900 | 900 | 900 | 900 |
| Minimum Frequency (mean entire) | 900 | 900 | 900 | 900 | 900 | 900 |
| Minimum Frequency(start) | 900 | 900 | 900 | 900 | 900 | 900 |
| Peak Frequency (end) | 900 | 900 | 900 | 900 | 900 | 900 |
| Peak Frequency (maximum) | 900 | 41000 | 15380 | 900 | 51700 | 13822.2 |
| Peak Frequency (mean) | 900 | 900 | 900 | 900 | 900 | 900 |
| Peak Frequency (Minimum entire) | 900 | 900 | 900 | 900 | 900 | 900 |
| Peak Frequency (start) | 900 | 900 | 900 | 900 | 900 | 900 |
| Peak Frequency (maximum entire) | 65400 | 69300 | 66380 | 1900 | 70300 | 44011.1 |
| Peak Frequency (mean entire) | 1000 | 1400 | 1080 | 900 | 900 | 900 |

##### 3.1.2.1. Fundamental Frequency and Harmonics

The sounds studied involved a sample of 1161, 1496 and 2342 calls for the female *A. gambiae*, male *A. gambiae* and *O. tormota* respectively. The parameters for the sounds of the male and female *A. gambiae* are given in Table 1. The calls for both male and female *A. gambiae* whose formants are given in the spectrogram given in Figure 3 reveal presence of the varied fundamental frequencies and their harmonics. The formants for the male *A. gambiae* appear weaker with some almost invisible. Harmonics are visible in the spectrograms of male and female *A. gambiae* stretching into ultrasonic levels. The degree of darkness of the spectrograms indicate that the acoustic energy of the female *A. gambiae* is greater than that of the male *A. gambiae.* The mean of the fundamental frequency (mean entire) was 1.00kHz and 955.56 Hz for the female and male *A. gambiae* respectively. The mean fundamental frequency (mean entire) of the sound samples of *O. tormota* was 4.89 kHz and 4.85 kHz higher than that of the male and female *A. gambiae* respectively. The sound of the male *A. gambiae* and *O. tormota* yielded a mean fundamental frequency of 900 Hz and 6.32 kHz respectively. The mean fundamental frequency (mean entire) of the female *A. gambiae* was higher than that of the male *A. gambiae.* Both sexes exhibited equal fundamental frequency (mean), Fundamental (mean), Fundamental (Minimum entire) and Fundamental (start) of 900Hz as given in Table 1. However, considerable variability was observed in the fundamental frequency (maximum entire) with the female recording the highest value of 3.66 kHz. A paired samples T-Test comparison of the fundamental frequency (maximum entire) of the sound of male *A. gambiae* by that of the sound of *O. tormota* at a confidence interval of 95% yielded the significance value, p = 0.001 < 0.05, implying that the fundamental frequencies differed significantly. There was a weak negative correlation (r = −0.130) between the fundamental frequency (maximum entire) of the sound of male *A. gambiae* and that of the sound of *O. tormota*.

The maximum frequency (minimum entire) of both sexes of *A. gambiae* was equal (1.90 kHz) with variability being observed in maximum frequency (end), maximum frequency (maximum), maximum frequency (mean), maximum frequency (maximum entire) and maximum frequency (mean entire). Frequency (maximum). A paired samples T-Test comparison of the maximum frequency (mean), maximum frequency (maximum), maximum frequency (end), maximum frequency (maximum entire) and maximum frequency (mean entire) of the sound of the female *A. gambiae* by that of the male *A. gambiae* at a significance level of 0.05 indicated no significant difference between the sounds of both sexes (p > 0.05). The maximum frequency (mean) for both sexes of *A. gambiae* correlated highly negative (r = −0.658). A low negative correlation was observed in the maximum frequency (maximum) (r = −0.041) and maximum frequency (end) (r = −0.291) for the sound for the sound of the male and female *A. gambiae.* The sound of the male and female *A. gambiae* correlated highly positive in maximum frequency(maximum entire) (r = 0.582) and maximum frequency (mean entire) (r = 0.517).

The minimum frequency (end), minimum frequency (maximum), minimum frequency (maximum entire), minimum frequency (mean), minimum frequency (mean entire), minimum frequency (minimum entire) and minimum frequency (start) for the sound of the male *A. gambiae* was 900 Hz which corresponded to its fundamental frequency. The minimum frequency (maximum entire) of the sound of the female *A. gambiae* exceeded that of the male *A. gambiae* by 1.00 kHz as given in Table 1. The peak frequency (end), peak frequency (mean), peak frequency (minimum entire) and peak frequency (start) for the sound of both sexes of the *A. gambiae* was 900 Hz. However, variation in peak frequency was observation in values for the peak frequency (maximum entire), Peak Frequency (mean entire) and Peak frequency (maximum). The mean of the Peak Frequency (maximum entire) for the female *A. gambiae* exceeded that of the male by 22.37 kHz. Comparing the sounds of the male and female *A. gambiae* showed no difference in Peak Frequency (maximum) (p = 0.280) and Peak Frequency (maximum entire) (p = 0.073) in a paired t-test comparison. A low negative correlation was observed in the Peak Frequency (maximum) (r = −0.429) whereas the Peak Frequency (maximum entire) yielded a high negative correlation (r = −0.722) for the comparison of the sounds of the male and female *A. gambiae*.

A significant difference was observed in the Peak Frequency (mean entire) (p = 0.01) in the paired sample t-test comparison of the sound of the male and female *A. gambiae.* The Peak Frequency (mean entire) of the sounds of the male and female *A. gambiae* correlated moderately positive (r = 0.536). The mean value of the peak frequency (mean entire) of the sound of the female *A. gambiae* was 180 Hz greater than that of the male *A. gambiae*. The maximum value of the maximum frequency (maximum entire) for sound of *O. tormota* was 4.90 kHz more than that of the male *A. gambiae.* However, the mean of the maximum frequency (maximum entire) for sound of *A. gambiae* was 48.33 kHz above that of *O. tormota* though both were ultrasonic ranges. Also the mean value of the minimum frequency (mean) and minimum frequency (mean entire) for the sound of *O. tormota* was greater than that of the male *A. gambiae* by 4.08 kHz and 5.00 kHz respectively. The paired samples T-Test comparison of the maximum frequency (mean entire) of the sound of the male *A. gambiae* by that of the sound of *O. tormota* at a confidence interval of 95% yielded the significance value, p = 0.129 > 0.05, implying that there did not exist a significant difference in the maximum frequency (mean entire) of the two sounds. A strong positive correlation at a Pearson’s product moment correlation coefficient, r = 0.679 existed between the maximum frequency (mean entire) of the sound of the male *A. gambiae* and that of the sound of *O. tormota.* Also a paired samples T-Test comparison of the maximum frequency (maximum entire) of the sound of the male *A. gambiae* by that of the sound of *O. tormota* at a confidence interval of 95% showed a significance difference in maximum frequency (maximum entire) (p = 0.011 < 0.05) for both sounds. However, there existed a weak positive correlation (r = 0.025) between the maximum frequency (maximum entire) of the sound of the male *A. gambiae* and that of the sound of *O. tormota.* The mean value of the peak frequency (end), peak frequency (mean), peak frequency (mean entire), peak frequency (minimum entire) and peak frequency (start) for the sound of *O. tormota* exceeded those of the sound of the male *A. gambiae* save for peak frequency (maximum entire) and peak frequency (maximum). The Peak Frequency (end) of the sound of the male *A. gambiae* and that of the sound of *O. tormota* differed significantly high (p = 0.001 < 0.05) with negligible positive correlation. It was also established that a high significance difference existed in the Peak Frequency (minimum entire), Peak Frequency (mean entire) and Peak Frequency (mean) in the paired t-test comparison (p <<< 0.05) with no correlation in the sound of the male *A. gambiae* and *O. tormota.* However, the Peak Frequency (maximum entire) (p = 0.020) and Peak Frequency (maximum) (p = 0.629) showed no significance difference in the comparison of the sound of the male *A. gambiae* and *O. tormota*. The Peak Frequency (maximum entire) (r = 0.329) and Peak Frequency (maximum) (r = 0.329) exhibited a low positive correlation. A low negative correlation in the sound of the male *A. gambiae* and *O. tormota* was noted in the Peak Frequency (maximum) (r = −0.404) with no significant difference (p = 0.629).

##### 3.1.2.2. Bandwidth

The sound of both the male and female *A. gambiae* yielded values of the minimum, maximum and mean Bandwidth (minimum entire) of 900 Hz. Variation in parameters of the sound of the male and female *A. gambiae* was observed in the bandwidth (end), bandwidth (maximum), bandwidth (maximum entire), bandwidth (mean) and bandwidth (mean entire). The female *A. gambiae* was characterised by a wide bandwidth compared to that of the male *A. gambiae* in terms of Bandwidth (end), Bandwidth (maximum entire), Bandwidth (mean) and Bandwidth (mean entire) as illustrated in Table 4. However, the mean of Bandwidth (maximum) was widest in the male *A. gambiae* compared to that of the female *A. gambiae* by 4.95 kHz. There existed a highly significant difference between bandwidth parameters of the sound of the male *A. gambiae* and that of *O. tormota* in terms of bandwidth (end) (p = 0.015 < 0.05), bandwidth (maximum entire) (p = 0.005< 0.05), Bandwidth (mean entire) (p = 0.027 < 0.05) and bandwidth (maximum) (p = 0.020 < 0.05) determined through a paired t-test comparison. However there did not exist a difference in bandwidth (mean) (p = 0.674 > 0.05) for the sound of the male *A. gambiae* and *O. tormota.* A positive correlation was observed in Bandwidth (end) (r = 0.536), bandwidth (maximum) (r = 0.896) and Bandwidth (mean entire) (r = 0.642) for the sound of the male *A. gambiae* and *O. tormota*. Uniquely, the two sounds exhibited a weak positive correlation in Bandwidth (mean) of 0.0560, bandwidth (maximum entire) (r = 0.005), and Bandwidth (mean) (r = 0.056). Table 4 gives the tabulated analysis of the Bandwidth parameters for the sounds of male and female *A. gambiae*. At 95% Confidence Interval of the difference between the bandwidth (end) of the sounds of the male and female *A. gambiae* was not significant (p = 0.5490 > 0.05) as established from the Paired Samples T Test. The bandwidth (end) of the sounds of the male and female *A. gambiae* correlated highly and positively (r = 0.6350). Also there did not exist any difference in bandwidth (maximum) between the sound of the male and female *A. gambiae* (p = 0.6340 > 0.05). The bandwidth (maximum) of the sound of the male and female *A. gambiae* recorded a very strong positive correlation (r = 0.9490).

**Table 2:**
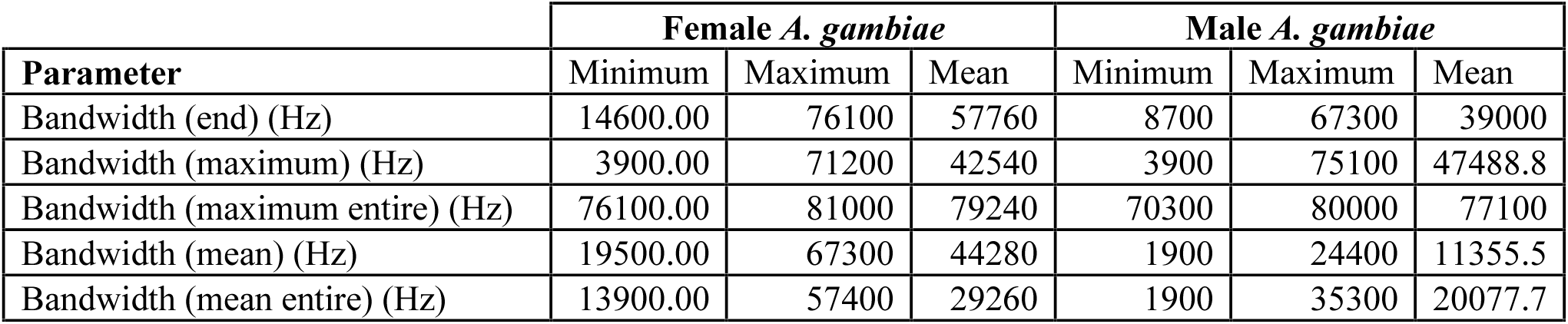
The Bandwidth parameters

| Parameter | Female *A. gambiae* |  |  | Male *A. gambiae* |  |  |
| --- | --- | --- | --- | --- | --- | --- |
|  | Minimum | Maximum | Mean | Minimum | Maximum | Mean |
| Bandwidth (end) (Hz) | 14600.00 | 76100 | 57760 | 8700 | 67300 | 39000 |
| Bandwidth (maximum) (Hz) | 3900.00 | 71200 | 42540 | 3900 | 75100 | 47488.8 |
| Bandwidth (maximum entire) (Hz) | 76100.00 | 81000 | 79240 | 70300 | 80000 | 77100 |
| Bandwidth (mean) (Hz) | 19500.00 | 67300 | 44280 | 1900 | 24400 | 11355.5 |
| Bandwidth (mean entire) (Hz) | 13900.00 | 57400 | 29260 | 1900 | 35300 | 20077.7 |

The bandwidth (maximum entire) and bandwidth (mean entire) for the sound of the male and female *A. gambiae* did not differ with p-values given as p = 0.0670 and 0.1610 respectively. The bandwidth (maximum entire) and bandwidth (mean entire) for the sound of both sexes of mosquitoes yielded a low positive correlation of r = 0.3010 and r = 0.3750 respectively. The bandwidth (mean) of the sound of the male and female *A. gambiae* differed significantly (p = 0.047 < 0.05) and correlated positively (r = 0.228).

##### 3.1.2.3. Amplitude

The study established statistical data for the amplitude parameters which included peak amplitude (end), peak amplitude (maximum), peak amplitude (maximum entire), peak amplitude (mean) and peak amplitude (mean entire) as given in Table 5. The peak amplitude (mean), Peak amplitude (mean entire) and peak amplitude (end) for the sound of the male *A. gambiae* were greater than that of the female *A. gambiae* by 0.9758 Hz and 0.9728 Hz respectively as given in Table 5. A paired sample T-test analysis at 0.05 confidence level showed that the there existed a high significance difference between the peak amplitude (mean) of the sound of the male *A. gambiae* and female *A. gambiae* (p = 0.001) with a high positive correlation (r = 0.5360). The same analysis on the peak amplitude (mean entire) of the sound of the male *A. gambiae* and female *A. gambiae* also revealed a significantly high difference (p = 0.002) with an equally high positive correlation (r = 0.534). However, there existed no difference between peak amplitude (end) of the sound of the male *A. gambiae* and female *A. gambiae* (p = 0.112) with a high negative correlation (r = −0.770). The peak amplitude (maximum) and peak amplitude (maximum entire) for the sound of the male and female *A. gambiae* were equal as given in Table 5. Also, the amplitude (maximum) and peak amplitude (maximum entire) of the sound of the male *A. gambiae* were equal but less than those of the female *A. gambiae* by 2.1821 Pa. Notably, there existed no difference in the peak amplitude (maximum) in the sound of the male *A. gambiae* and female *A. gambiae* (p = 0.089) with a low positive correlation (r = 0.157). The difference and trend in peak amplitude (maximum entire) in the sound of the male *A. gambiae* and female *A. gambiae* was equal to that in peak amplitude (maximum). A paired t-test comparison of the Peak amplitude (maximum), peak amplitude (mean), Peak amplitude (mean entire) and peak amplitude (maximum entire) of the sound of the male *A. gambiae* by the sound of *O. tormota* showed a highly significant difference (p << 0.05). There existed a low positive correlation in peak amplitude (mean) (r = 0.479) and peak amplitude (mean entire) (r = 0.477) for the sound of the male *A. gambiae* and *O. tormota*. However, a low negative correlation (r = −0.004) was observed in the comparison of the peak amplitude (maximum) and peak amplitude (maximum entire) for the sound of the male *A. gambiae* by the sound of *O. tormota*.

**Table 3:** The Amplitude parameters

| Parameter (in Pa) | Female *A. gambiae* |  |  | Male *A. gambiae* |  |  |
| --- | --- | --- | --- | --- | --- | --- |
|  | Minimum | Maximum | Mean | Minimum | Maximum | Mean |
| Peak amplitude (end) | 81.79 | 89.4 | 85.3679 | 87.19 | 89.25 | 88.1488 |
| Peak amplitude (maximum) | 93.48 | 96.97 | 95.132 | 90.68 | 95.6699 | 92.9499 |
| Peak amplitude (maximum entire) | 93.48 | 96.97 | 95.132 | 90.68 | 95.6699 | 92.9499 |
| Peak amplitude (mean) | 87.5 | 88.37 | 87.8819 | 88.57 | 89.1399 | 88.8577 |
| Peak amplitude (mean entire) | 87.5 | 88.3899 | 87.886 | 88.57 | 89.1399 | 88.8588 |
| Peak amplitude (start) | 40.65 | 45.5099 | 43.7339 | 41.5499 | 43.49 | 43.0588 |

##### 3.1.2.4. Acoustic Energy and Power

The acoustic energy of the male *A. gambiae* was greater than that of *O. tormota*. The sound of male *A. gambiae* possessed a mean of the acoustic energy of 82.3197 Pa^2^s which was six times the acoustic energy in *O. tormota*. The maximum acoustic energy in the entire sound spectrum of the sound of male *A. gambiae* was 140.419 Pa^2^s, 3.13 times greater than that of the sound of *O. tormota*. The sound of *O. tormota* yielded the least minimum acoustic energy of 0.00097 Pa^2^s which was 25,639.6 times less than the acoustic energy of the sound of male *A. gambiae*. A paired sample T-test analysis of the acoustic energy for the male *A. gambiae* by that of the *O. tormota* at a significance level of 0.05 yielded a significant difference between the two sounds (p = 0.013) which correlated negatively high (r = 0.630). The signal power within the 10-90 kHz range of study for the sounds of male *A. gambiae*, male *A. gambiae* and *O. tormota* are given in Figure 5, 6 and 7 respectively. The signal power for the sounds of the male *A. gambiae* remained almost constant at 80 dB from 10 kHz to 65 kHz beyond which the acoustic energy declined to 45 dB. The sound was not pulsate. The sounds of the female *A. gambiae* did not exhibit any spikes in power but remained steady at 85 dB from 10 kHz up to 60 kHz beyond which the acoustic energy declined to 50 dB. The signal for the female *A. gambiae* was not pulsate though possessed higher acoustic power compared to the signal of the male *A. gambiae*. The signal power spectrum for *O. tormota* was pulsate due to characteristic spikes. The acoustic power was less than that of both sexes of the *A. gambiae*. The peak acoustic power (maximum entire) was 89 dB with the acoustic power (minimum entire) being 41 dB, maintaining an average acoustic power of 55 dB. The acoustic power (minimum entire) was maintained at 41 dB up to 90 kHz.

**Figure 5:**
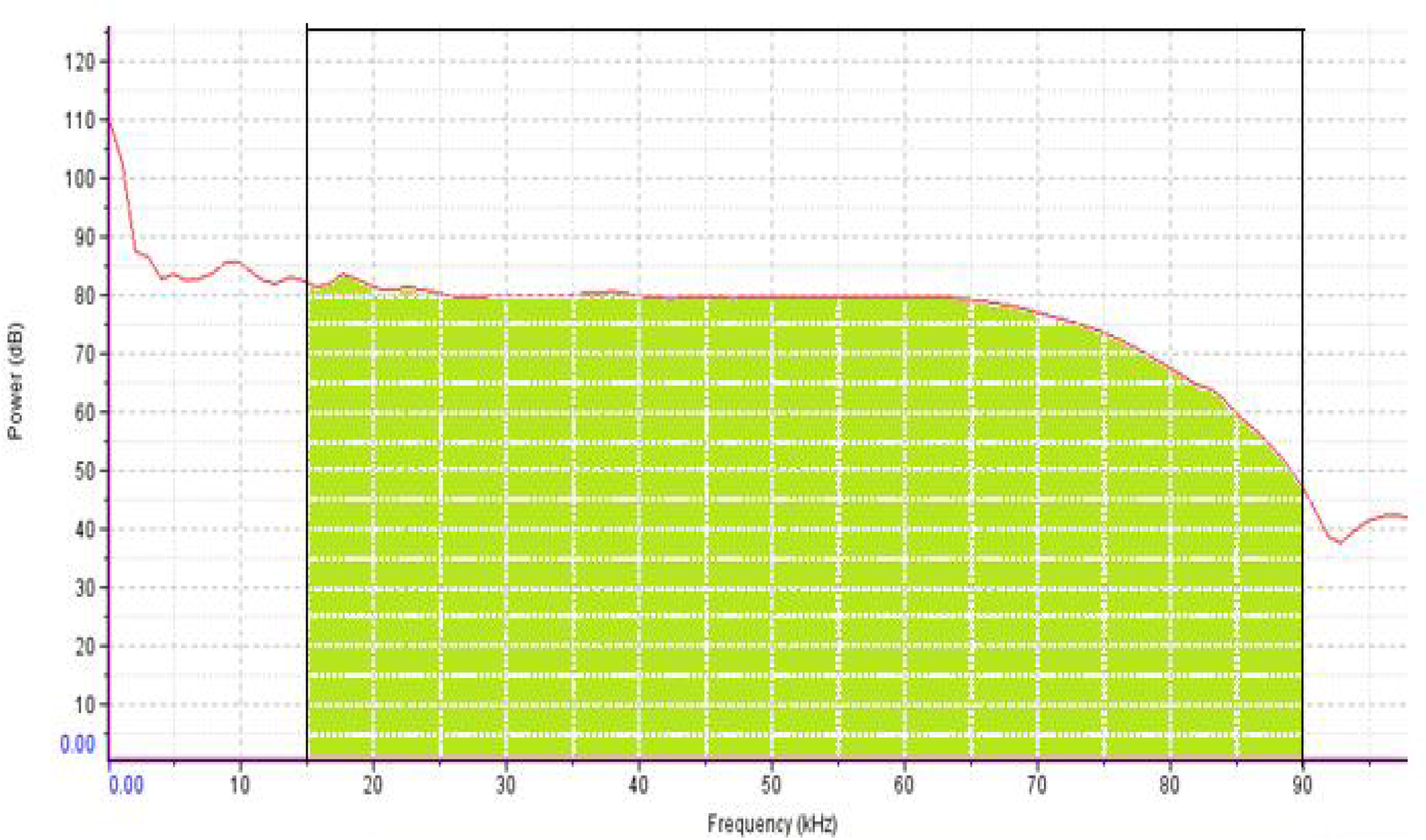
Signal power spectrum of the sound of the male *A. gambiae*

**Figure 6:**
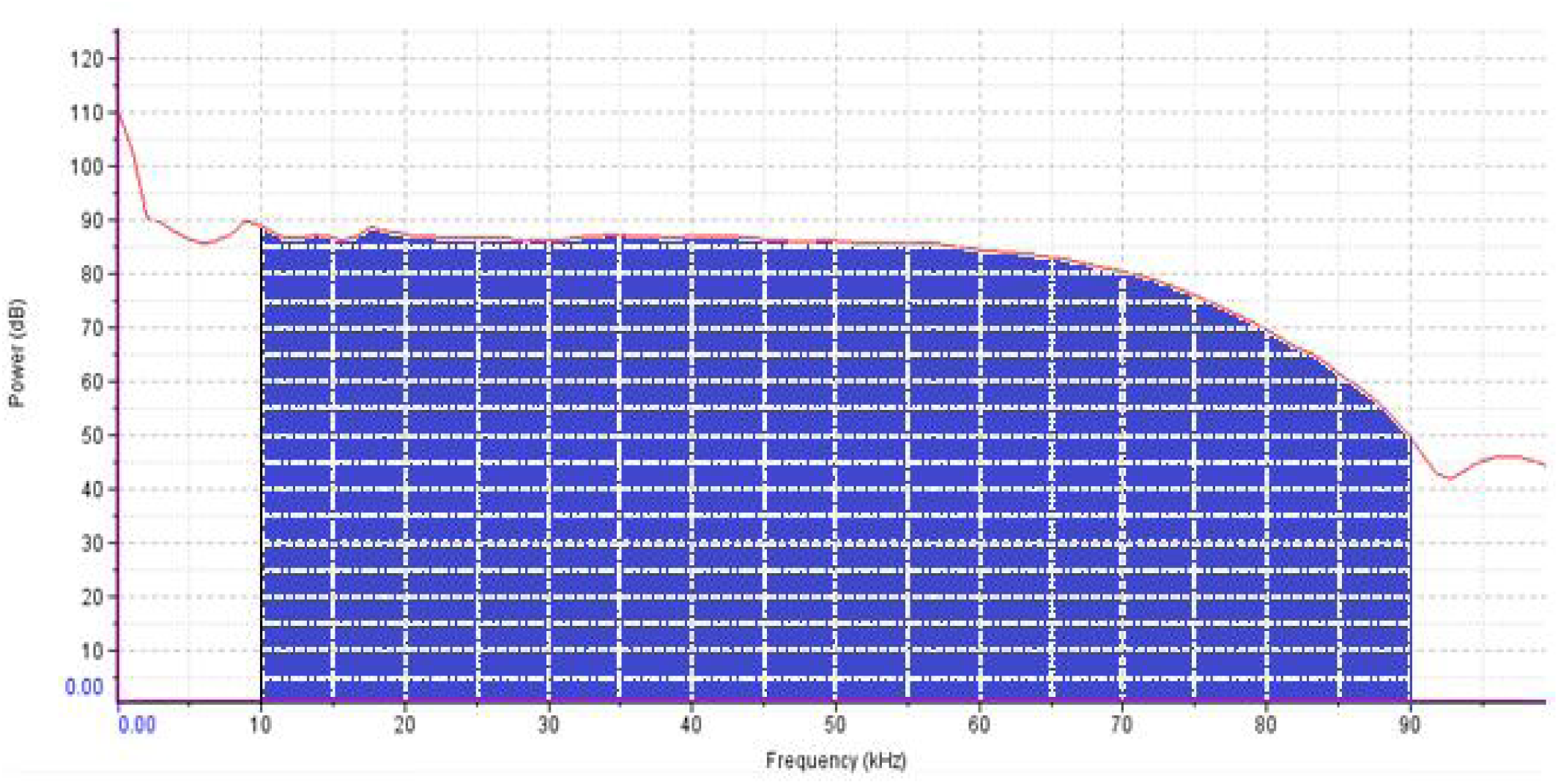
Signal power spectrum of the sound of the female *A. gambiae*

**Figure 7:**
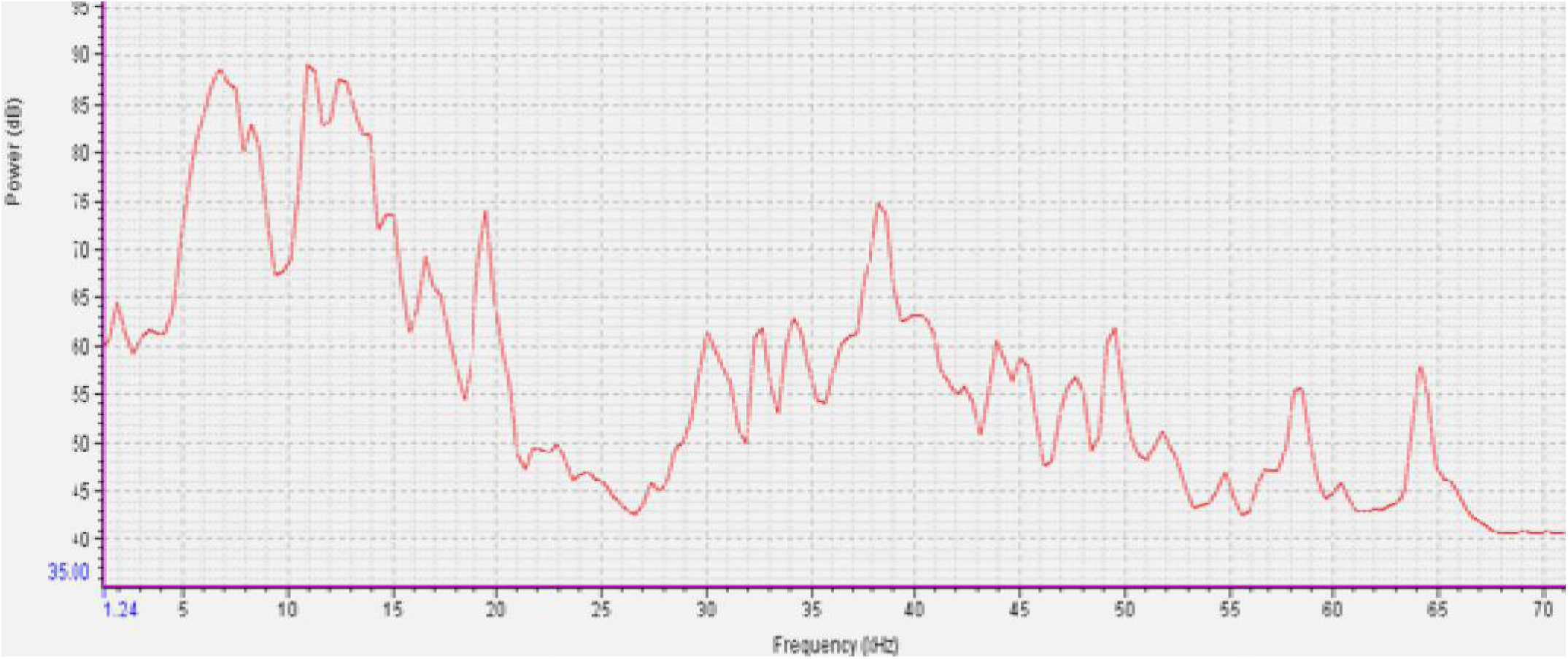
Signal power spectrum of the sound of *O. tormota*

## Conclusion

The male and female *A. gambiae* generate sounds through wing-beat whereas *O. tormota* produces the sound through vocal apparati. The spectral features and formants in the sounds of the male and female *A. gambiae*, and *O. tormota* revealed presence of frequency modulation (FM), varied fundamental frequencies and their harmonics. The *O. tormota* formants also showed constant frequency (CF). All sounds consisted of several harmonic segments with energy extending into the ultrasonic range, at some instances, exhibiting signal breaks, subharmonics and a chaotic nonlinear characteristics. The sound of *O. tormota* was also characterised by constant frequency (CF) modulation. The acoustic energy of the female *A. gambiae* was greater than that of the male *A. gambiae* with mean of the fundamental frequency (mean entire) of 1.00kHz and 955.56 Hz respectively. Both sexes showed equal fundamental frequency (mean), Fundamental (mean), fundamental (Minimum entire), Bandwidth (minimum entire) and Fundamental (start) of 900Hz and maximum frequency (minimum entire) of 1.90 kHz. The peak amplitude (maximum) and peak amplitude (maximum entire) for both sexes of the *A. gambiae* was also equal. However, the male *A. gambiae* recorded the highest peak amplitude (mean), peak amplitude (mean entire) and peak amplitude (end) compared to the female *A. gambiae*. The sounds of male *A. gambiae* and *O. tormota* differed significantly high in the fundamental frequency (maximum entire), Peak amplitude (maximum), peak amplitude (mean), Peak amplitude (mean entire) and peak amplitude (maximum entire). In a paired t-test comparison of the sounds of the male *A. gambiae* and *O. tormota*, the differences in the maximum frequency (maximum entire), bandwidth (end), bandwidth (maximum) and bandwidth (mean entire) was highly significant. However, there existed no significant difference in the maximum frequency (mean entire) for the sounds of the male *A. gambiae* and *O. tormota*. The minimum frequency (maximum), minimum frequency (mean), minimum frequency (minimum entire), minimum frequency (mean entire) and minimum frequency (start) for both the male and female *A. gambiae* was equal to 900 Hz. However, the peak Frequency (maximum) of the sound of male *A. gambiae* compared by the sound of *O. tormota* showed no significant differences (p = 0.629 > 0.05). The sound of the male and female *A. gambiae* did not differ significantly (0.280) with a low negative correlation (r = −0.429). There existed no significant difference in bandwidth (end), bandwidth (maximum), bandwidth (maximum entire), peak amplitude (end), peak amplitude (maximum) and bandwidth (mean entire) of the sound of the male and female *A. gambiae* with the exception of the acoustic energy, peak amplitude (mean), peak amplitude (mean entire) and bandwidth (mean). The acoustic energy of the sound male *A. gambiae* was greater than that of *O. tormota*, but less than that of the sound of female *A. gambiae.* The maximum signal power in *O. tormota*, female *A. gambiae* and male *A. gambiae* was 89 dB, 85 dB and 80 dB respectively. These parameters of the sound male *A. gambiae* and *O. tormota* compare favourably hence the need for further investigation on the sound of the male *A. gambiae* on its feasibility to startle the female *A. gambiae*.

## Acknowledgement

We thank Masinde Muliro University of Science and Technology and Egerton University for their immense support during the research. Our sincere gratitude is extended to our colleagues in the Department of Physics of Egerton University, Moi University and Masinde Muliro University of Science and Technology. We are highly indebted to Prof. Feng, Dr. Ndinya, Raimund Specht of Avisoft Bioacoustics, Pettersson Elektronik AB and Cornell Lab of Ornithology for their kind donations and support. We also thank the Staff in KEMRI (Kisumu) for their encouragement and valuable input.

## References

Abdolali, A., Hasanzade, H. and Salary, M. M. (2013). The antenna analysis of insect antennae. World Journal of Modelling and Simulation. 9 (3): 235–240

Andrade, C. F. S. and Bueno, V. S. (2001). Evaluation of Electronic Mosquito-Repelling Devices Using *Aedes albopictus* (Skuse) (Diptera: Culicidae). Neotropical Entomology. 30(3): 497–499

Antonelli, A., Murray, T and Danies, C. (2007). Pest Management for Prevention and Control of Mosquitoes with Special Attention to West Nile Virus. WSU-Puyallup.

Arch, V. S., Grafe T. U. and Narins, P. M. (2008). Ultrasonic signalling by a Bornean frog. Biology Lett. 4: 19–22.

Arthur, B. J., Emr, K. S., Wyttenbach, R. A. and Hoyd, R. R. (2014). Mosquito (Aedes aegypti) flight tones: Frequency, harmonicity, spherical spreading, and phase relationships. The Journal of the Acoustical Society of America. 135 (2): 933–941.

Ashim, A. K., Tanveerul, H., Shaik, N., Farzand, A. and Parrmeshear, G. (2017). Design Considerations of Mosquito repellent unit. International Journal of Innovative Research in Technology. 3(9): 20–23

Baldini, F., Gabrieli, P., South, A., Valim, C., Mancini, F. and Catteruccia, F. (2013). “The interaction between a sexually transferred steroid hormone and a female protein regulates oogenesis in the malaria mosquito *Anopheles gambiae*,” PLOS Biology, 11:e1001695, 2013. https://doi.org/10.1371/journal.pbio.1001695

CDC. (2010). *Anopheles* mosquitoes. Malaria. Centers for Disease Control and Prevention 1600 Clifton Rd. Atlanta, GA 30329-4027, USA 800- CDC-INFO (800-232-4636) TTY.

Clements, A. N. (1992). The Biology of Mosquitoes: Development, Nutrition and Reproduction. Chapman & Hall, London.

Diabaté, A., Yaro, A. S., Dao, A., Diallo, M., Huestisa, D. L. and Lehmann, T. (2011). Spatial distribution and male mating success of *Anopheles gambiae* swarms. BMC Evolutionary Biology. 11: 184.

Enayati, A., Hemingway, J. and Garner, P. (2010). Electronic mosquito repellents for preventing mosquito bites and malaria infection (Review). The Cochrane Library Journal. 3: 1 – 16.

Fei, L., Ye, C.-Y., Huang, Y.-A. and Liu, M.-Y. 1999. Atlas of Amphibians of China. Henan Science and Technical Press, Zhengzhou.

Feng, S. A., Riede, T., Arch, V. S., Yu, Z., Xu, Z., Yu, X. and Shen, J. (2009). Diversity of the Vocal Signals of Concave-Eared Torrent Frogs (*Odorrana tormota*): Evidence for Individual Signatures. International Journal of Behavioural Biology. doi: 10.1111/j.1439-0310.2009.01692.x: 1–15

Feng, A. S., Narins, P. M., Xu, C. H., Lin, W.Y and Yu, Z. L. (2006). Ultrasonic communication in frogs. Nature. 440: 333–336.

Feng, A. S., Narins, P. M. and Xu, C. H. (2002). Vocal acrobatics in a Chinese frog, *Amolops tormotus*. Naturwissenshaften Journal. 89: 352–356.

Foster, W. A. and Walker, E. D. (2009). *Mosquitoes* (Culicidae). In Mullen, G, Durden, L. (Eds.) Medical and veterinary entomology. 2nd Ed. Academic Press, Burlington, MA. 637: 207–259

Frost, D.R. 2013. Amphibian Species of the World: an Online Reference. Version 5.6 (9 January 2013). Electronic Database. American Museum of Natural History, New York, USA. Available at: http://research.amnh.org/herpetology/amphibia/index.html.

Garros, C., Ngugi, N., Githeko, A. E., Tuno, N., & Yan, G. (2008). Gut content identification of larvae of the Anopheles gambiae complex in western Kenya using a barcoding approach. Molecular ecology resources, 8(3): 512–8.

Gibson, G., Warren, B., and Russell, I. J. (2010). Humming in Tune: Sex and Species Recognition by Mosquitoes on the Wing. Journal of the Association for Research in Otolaryngology. 11: 527–540.

Gibson, G. and Russell, I. (2006). Flying in tune: sexual recognition in mosquitoes. Current Biology. 16: 1311–1316.

Gillies, M. T. and de Meillon, B. (1968). The *Anophelinae* of Africa south of the Sahara (Ethiopian Zoogeographical Region). Publications of the South African Institute for Medical Research. 54: 1–343.

Göpfert, M. C. and Robert, D. (2000). Nanometre-range acoustic sensitivity in male and female mosquitoes. Proceedings of the Royal Society of London B: Biological Sciences. 267: 453–457

Göpfert, M. C., Biegel, H. and Robert, D. (1999) Mosquito hearing: sound-induced antennal vibrations in male and female *Aedes aegypti*. The Journal of Experimental Biology. 202: 2727–2738

Huiqing, G. and Ermi, Z. (2004). Odorrana tormota. The IUCN Red List of Threatened Species.2004: e.T58226A11752893. http://dx.doi.org/10.2305/IUCN.UK.2004.RLTS.T58226A11752893.en

Hoy R. (2006). A boost for hearing in mosquitoes. Proceedings of the National Academy of Sciences of the United States of America, 103(45), 16619–20.

Hoy, R. R. and Robert, D. (1996). Tympanal hearing in insects. Annual Revews of Entomology. 41: 433–450.

Imam, H., Zarnigar, Sofi, G., & Seikh, A. (2014). The basic rules and methods of mosquito rearing (*Aedes aegypti*). Tropical parasitology. 4 (1): 53–5.

Kweka, E. J., Zhou, G., Munga, S., Lee, M. C., Atieli, H. E., Nyindo, M., Githeko, A. K. and Yan, G. (2012). *Anopheline* larval habitats seasonality and species distribution: a prerequisite for effective targeted larval habitats control programmes. Public Library of Science One. 7: 1–10.

Lapshin, D.N. and Vorontsov, D.D. (2018). Low-Frequency Sounds Repel Male Mosquitoes *Aedes diantaeus* N.D.K. (*Diptera, Culicidae*). Entomologicheskoe Obozrenie. 97(2): 194–202.

Mang’are, P. A., Maweu, O. M., Ndiritu, F. G. and Vulule, J. M. (2012). Determination of Acoustic Transmission Parameters of the Sound of *C. afra* and *A. tormotus*. International Journal of Biophysics. 2(4): 53–67

Maweu, O. M., Deng, A. L. and Muia, L. M. (2009).A Comparative study of *A. gambiae* male mosquito’s response to frequency modulated [FM] and pulse modulated [PM] waves at different acoustic frequencies and distances. Indonesian Journal of Physics. 20: 81–84.

Mohankumar, D. (2010). Ultrasound and insects. Electronics and Animal Science. https://dmohankumar.wordpress.com/2010/04/08/ultrasound-and-insects/. Accessed on 1-November-17 10:00 PM.

Nijhout, H. F. and Craig. G. B. (1971). Reproductive isolation in Stegomyia mosquitoes. III. Evidence for a sexual pheromone. Entomol. Exp. Appl. 14: 399–412.

Pennetier, C., Warren, B., Dabire, K. R., Russel, I. J. and Gibson, G. (2010). “Singing on the wing” as a mechanism for species recognition in the malarial mosquito *Anopheles gambiae*. Current Biology. 20(2): 131–136.

Raman, D. R., Gerhardt, R. R. and Wilkerson, J. B. (2007). Detecting insect flight sounds in the field: Implications for acoustical counting of mosquitoes. Trans ASABE 50(4):1481

Sane, P. S. and McHenry, M. J. (2009). The biomechanics of sensory organs. Integrative and Comparative Biology Advance Access.

Shen, J., Xu, Z., Yu, Z., Wang, S., Zheng, D and Fan, S. (2011). Ultrasonic frogs show extraordinary sex differences in auditory frequency sensitivity. Nature Communications. 2: 342.

Wishart G., and Riordan D. F. (1959). “ Flight responses to various sounds by adult males of *Aedes aegypti* (L.) (Diptera: Culicidae).” The Canadian entomologist. 91: 181–191. 10.4039/Ent91181-3

Wu, G.-f. 1977. A new species of frog from Huang-Shan, Anhui—*Rana tormotus* Wu. Acta Zoologica Sinica/ Dong wu xue bao. Beijing 23: 113–115.

